# A Neural Locus for Perceptually-Relevant Saccadic Suppression in Visual-Motor Neurons of the Primate Superior Colliculus

**DOI:** 10.1101/079384

**Authors:** Chih-Yang Chen, Ziad M. Hafed

**Affiliations:** Werner Reichardt Centre for Integrative Neuroscience, Tuebingen University, Tuebingen, BW, 72076, Germany; Graduate School of Neural and Behavioural Sciences, International Max Planck Research School, Tuebingen University, Tuebingen, BW, 72074, Germany

**Author notes:** **Corresponding Author:** Ziad M. Hafed Werner Reichardt Centre for Integrative Neuroscience Otfried-Mueller Str. 25 Tuebingen, BW, 72076, Germany.

**Keywords:** Saccades, Microsaccades, Superior colliculus, Saccadic suppression, Perceptual stability

## Abstract

Saccadic eye movements cause rapid retinal-image shifts that go perceptually unnoticed several times per second. The mechanisms for perceptual saccadic suppression have been controversial, in part due to sparse understanding of neural substrates. Here we uncovered an unexpectedly specific neural locus for saccadic suppression in the primate superior colliculus (SC). We first developed a sensitive behavioral measure of perceptual suppression in two male macaque monkeys (*Macaca mulatta*), demonstrating known selectivity to low spatial frequencies. We then investigated visual responses in either purely visual SC neurons or anatomically-deeper visual-motor neurons, which are also involved in saccade generation commands. Surprisingly, visual-motor neurons showed the strongest visual suppression, and the suppression was dependent on spatial frequency like in perception. Most importantly, visual-motor neuron suppression selectivity was highly predictive of behavioral suppression effects in each individual animal, with our recorded population explaining up to ~74% of behavioral variance even on completely different experimental sessions. In contrast, purely visual SC neurons only had mild and unselective suppression (only explaining up to ~48% of behavioral variance). These results run contrary to a hypothesized SC mechanism for saccadic suppression, in which a motor command in the visual-motor and motor neurons is relayed to the more superficial purely visual neurons to suppress them, and to then potentially be fed back to cortex. Instead, our results indicate that an extra-retinal modulatory signal mediating perceptual suppression is already established in visual-motor neurons.

**New & Noteworthy:** Saccades, which repeatedly re-align the line of sight, introduce spurious signals in retinal images that normally go unnoticed. In part, this happens because of peri-saccadic suppression of visual sensitivity. Here we discovered that a specific sub-type of superior colliculus (SC) neurons may play a critical role in saccadic suppression. Curiously, it is the neurons that help mediate the saccadic command itself that exhibit perceptually-relevant changes in *visual* sensitivity, not the previously hypothesized purely visual neurons.

## Introduction

A long standing question in visual neuroscience has been on the sense of perceptual stability that we normally experience despite incessant eye movement (Wurtz 2008). Saccadic eye movements dramatically alter retinal images several times per second. During each saccade, retinal images undergo rapid motion, which can be beyond the range of motion sensitivity of many neurons. Such motion ought, at least in principle, to cause a brief period of “grey out” every time a saccade occurs (Campbell and Wurtz 1978; Matin 1974; Wurtz 2008; Wurtz et al. 2011), much like the grey out experienced while standing near train tracks and high-speed trains sweep by.

Several fiercely debated theories on why we do not experience saccade-related visual disruptions have emerged. On the one hand, purely visual mechanisms, such as masking (Matin et al. 1972), can be sufficient to suppress perception of saccade-induced grey out and/or motion (Wurtz 2008). Consistent with this, people are not entirely “blind” during saccades, as long as spatio-temporal properties of peri-saccadic stimuli remain within sensitivity ranges of visual neurons (Burr and Ross 1982; Castet et al. 2001; Castet and Masson 2000; Garcia-Perez and Peli 2011; Ilg and Hoffmann 1993; Matin et al. 1972; Ross et al. 1996). On the other hand, extra-retinal mechanisms (Sperry 1950; von Holst and Mittelstaedt 1950) for perceptual suppression are supported by the lack of suppression during simulated image displacements (Diamond et al. 2000), the dependence of suppression on spatial frequency (Burr et al. 1982; Burr et al. 1994; Hass and Horwitz 2011; Volkmann et al. 1978), and the observation of saccade-related modulation of neural excitability in the absence of visual stimulation (Rajkai et al. 2008).

While it is likely that a combination of visual and extra-retinal mechanisms co-exist (Wurtz 2008), controversies surrounding saccadic suppression will not be resolved without further understanding of neural mechanisms. We were particularly interested in potential mechanisms for extra-retinal suppression, whose sources remain elusive. For example, it was suggested from behavioral studies that selective perceptual suppression of low spatial frequencies is evidence for selective magno-cellular (achromatic) pathway suppression (Burr et al. 1994). However, in lateral geniculate nucleus (LGN) and primary visual cortex (V1), the two earliest visual areas, selective magno-cellular suppression is not established (Hass and Horwitz 2011; Kleiser et al. 2004; Ramcharan et al. 2001; Reppas et al. 2002; Royal et al. 2006). In addition, a popular hypothesis about a source of saccadic suppression is that a “corollary” of saccade commands in visual-motor and motor neurons of the superior colliculus (SC) is fed back to superficial purely visual neurons to suppress their sensitivity, and to jumpstart a putative feedback pathway for cortical suppression through pulvinar (Berman and Wurtz 2008; 2010; 2011; Isa and Hall 2009; Lee et al. 2007; Phongphanphanee et al. 2011; Wurtz 2008; Wurtz et al. 2011). However, evidence for an SC saccadic suppression pathway from visual-motor/motor neurons to visual neurons comes primarily from rodent SC slices (Isa and Hall 2009; Lee et al. 2007; Phongphanphanee et al. 2011). In the awake, behaving primate, findings of stronger suppression in visual-motor rather than visual neurons cast doubt on this hypothesis (Chen et al. 2015; Hafed et al. 2015; Hafed and Krauzlis 2010).

We visited the question of neural loci for saccadic suppression by explicitly testing the predictions of a perceptually-relevant suppressive pathway from SC motor-related neurons to purely visual ones. We have previously shown that SC neurons exhibit time courses of saccadic suppression remarkably similar to those of perceptual effects (Hafed and Krauzlis 2010). Here, we adapted our sensitive behavioral paradigm (Hafed and Krauzlis 2010) to first establish *selectivity* in saccadic suppression, and we then asked whether *visual* neural modulations in either purely visual or visual-motor SC neurons would reflect such selectivity. Contrary to the hypothesis about a suppressive pathway from deep to superficial layers (Isa and Hall 2009; Lee et al. 2007; Phongphanphanee et al. 2011), we observed perceptually-relevant and spatial-frequency specific saccadic suppression only in the deeper visual-motor neurons. Our results suggest that the SC is indeed relevant for perceptually-relevant saccadic suppression, but that the putatively extra-retinal modulatory signal mediating suppression may already be established in the visual-motor neurons, without the need for a relay through purely visual neurons.

## Materials and Methods

We exploited microsaccades to study saccadic suppression because they offer important experimental advantages while at the same time being mechanistically similar to larger saccades (Hafed 2011; Hafed et al. 2015; Hafed et al. 2009; Zuber et al. 1965). First, microsaccades are small (median amplitude in our data: ~7.5 min arc). Thus, pre- and post-movement visual response fields (RF’s) are not displaced by much, minimizing the problem of dramatic spatial image shifts caused by saccades (Wurtz 2008; Wurtz et al. 2011). Experimentally, this meant presenting the exact same stimulus at the exact same screen location with and without microsaccades to isolate suppression effects. Second, microsaccades have velocities significantly <100 deg/s (median peak velocity in our data: ~17.7 deg/s). Thus, image motion caused by microsaccades is well within the range of motion sensitivity, even for small features (Thiele et al. 2002), allowing us to study suppression even when no motion-induced grey out is expected to occur. Third, we have previously shown that SC visual sensitivity exhibits pre-, peri-, and post-microsaccadic suppression that is very similar in time course and amplitude to perceptual saccadic suppression in humans, and we have also demonstrated a sensitive behavioral paradigm for the same phenomenon (Hafed and Krauzlis 2010). Fourth, and most importantly, we avoided potential masking effects by only presenting stimuli immediately *after* microsaccades. This allowed us to study a potential intrinsic reduction in neural excitability after saccades (Chen et al. 2015; Hafed and Krauzlis 2010; Zuber et al. 1966), and to ensure comparing “no-microsaccade” to “microsaccade” conditions without the latter involving saccade-induced retinal image motion. Thus, the logic of all of our experiments was to present high-contrast gratings (80% contrast), which were highly visible and well within the saturation regime of SC contrast sensitivity curves (Chen et al. 2015; Hafed and Chen 2016; Li and Basso 2008), and to ask whether either behavioral or visual neural responses to these gratings were altered if the gratings appeared immediately after a microsaccade.

### Animal Preparation

Ethics committees at regional governmental offices in Tuebingen approved experiments. Monkeys P and N (male, *Macaca mulatta*, aged 7 years) were prepared earlier (Chen and Hafed 2013; Chen et al. 2015; Hafed and Chen 2016; Hafed and Ignashchenkova 2013). We used scleral search coils to record eye movements (Fuchs and Robinson 1966; Judge et al. 1980).

### Behavioral Tasks

In all tasks, monkeys initially fixated a small, white spot presented over a gray background (Chen and Hafed 2013; Chen et al. 2015; Hafed and Ignashchenkova 2013).

#### Behavioral tests

Trials started with an initial fixation interval of random duration (between 600 and 1500 ms). After this interval, we initiated a real-time process to detect microsaccades (Chen and Hafed 2013). If a microsaccade was detected within 500 ms, a stationary vertical Gabor grating appeared at 3.5 deg to the right or left of fixation, and the fixation spot was removed simultaneously. Monkeys oriented to the grating as fast as possible using a saccadic eye movement, and saccadic reaction time (RT) served as a sensitive behavioral measure of SC visual response strength (Boehnke and Munoz 2008; Hafed and Chen 2016; Hafed et al. 2015; Hafed and Krauzlis 2010; Tian et al. 2016).

Grating onset occurred ~25, 50, 75, 100, 150, or 200 ms after online microsaccade detection, and we later measured precise times of microsaccade onset during data analysis for all results presented in this paper (see Data Analysis below). If no microsaccade was detected during our 500 ms online detection window, a grating was presented anyway, and the data contributed to “baseline” measurements (i.e. ones with the stimulus appearing without any nearby microsaccades). The grating was 2 deg in diameter. Spatial frequency in cycles per degree (cpd) was one of five values: 0.56, 1.11, 2.22, 4.44, 11.11 (Hafed and Chen 2016), and phase was randomized. Our monitor resolution allowed displaying the highest spatial frequency without aliasing and distortion. We collected 8153 and 7117 trials from monkeys N and P, respectively. We removed trials with an intervening microsaccade between fixation spot removal and the orienting saccade.

#### Neural recordings

We isolated single neurons online, and we identified their RF locations and sizes using standard saccade tasks (Chen et al. 2015; Hafed and Chen 2016). We then ran our main experimental paradigm. In each trial, monkeys fixated while we presented a similar vertical grating to the one we used in behavioral tests above, but the grating was now inside the recorded neuron’s RF. Grating size was optimized for the recorded neuron, and was specifically chosen to fill as much of the RF as possible (and showing >1 cycle of the lowest spatial frequency). Task timing was identical to that in (Chen et al. 2015); briefly, a grating was presented for 250 ms while monkeys fixated, and the monkeys never generated any saccadic or manual responses to the grating (they simply maintained fixation, during which they generated microsaccades, and they were rewarded at the end of the 250 ms stimulus presentation phase for maintaining fixation). We collected data from 90 neurons, covering 1-24 deg eccentricities. We classified neurons as purely visual neurons or visual-motor neurons using previous criteria from visually-guided and memory-guided saccade tasks (Chen et al. 2015; Hafed and Chen 2016). To ensure sufficient microsaccades for statistical analyses (i.e. with sufficient trials having stimulus onset within the critical post-movement intervals that we analyzed), we collected >800 trials per neuron. We then separated trials as ones having no microsaccades within +/- 100 ms from grating onset (>100 trials per neuron; mean: 289 trials per neuron) or ones with grating onset within 50 ms *after* microsaccades (>25 trials per neuron; mean: 79 trials per neuron). For some analyses, we considered grating onsets up to 100 ms after microsaccades.

It is important to note here that for all neurons reported in this paper, we never observed a microsaccade-related movement burst (Hafed et al. 2009; Hafed and Krauzlis 2012).

Thus, even for stimuli appearing immediately after a microsaccade, the neural responses that we analyzed were *visual* bursts in response to stimulus onset, and not movement-related saccade or microsaccade bursts. The only difference between purely visual and visual-motor neurons in this study was that visual-motor neurons would, in principle, exhibit a saccade-related burst if the monkeys were to hypothetically generate saccades towards the RF location (but not if they generated smaller microsaccades during fixation). Thus, any neural modulations that we report in this study are not direct microsaccade-related motor bursts. It is also important to note that our monkeys did not generate any targeting saccades to the gratings during recordings. We were simply studying *visual* sensitivity if a stimulus appeared near an eye movement (which happened to be small due to our experimental choice). Our approach was thus similar to classic ways of studying neural correlates of saccadic suppression (i.e. monkeys make saccades while neurons are visually stimulated; e.g. Zanos et al. 2016; Hafed and Krauzlis 2010; Bremmer et al. 2009).

### Data Analysis

#### Behavioral analyses

For behavior, we measured RT as a function of spatial frequency and time of grating onset relative to microsaccades. We detected microsaccades using previously described methods (Hafed et al. 2009), and we used such detection to identify grating onset time relative to microsaccade onset or offset. We defined no microsaccade trials as trials with no microsaccades <250 ms from grating onset. RT on these trials constituted our baseline.

#### Firing rate analyses

For neural data, we measured stimulus-evoked firing rate after the onset of a given spatial-frequency grating under two scenarios: (1) when the grating appeared without any nearby microsaccades within +/- 100 ms, and (2) when the grating appeared immediately after a microsaccade. Baseline, no-microsaccade spatial frequency tuning curves (i.e. responses for each given spatial frequency) were described recently (Hafed and Chen 2016), but here we analyzed microsaccadic influences on these curves. We did not analyze trials with grating onset immediately before or during microsaccades, to avoid pre-movement modulations (Chen et al. 2015; Hafed 2013) and retinal-image shift effects caused by movement of the eyes.

To analyze stimulus-evoked firing rate, we measured peak visual response 20-150 ms after grating onset. In some analyses, we also measured mean firing rate within an interval after grating onset. In this case, we tailored the averaging window to each spatial frequency, because visual burst latency depended on spatial frequency (e.g. Fig. 3, see cyan bars on x-axes). Our averaging windows for mean firing rate were all 100-ms long windows starting at 20 ms (for 0.56 cpd), 25 ms (for 1.11 cpd), 30 ms (for 2.22 cpd), 40 ms (for 4.44 cpd), or 80 ms (for 11.11 cpd). We chose these intervals after inspecting all neurons as follows. For each neuron and spatial frequency, we found the time at which firing rate became >3 s.d. from a pre-stimulus baseline (mean activity 0-100 ms before stimulus onset). We then chose our window start time as the average start time of the first quartile of the entire population (rounded to the nearest 5 ms).

To compare visual sensitivity on microsaccade and no-microsaccade trials, we created a “normalized firing rate” modulation index for each individual spatial frequency. We measured firing rate on microsaccade trials (i.e. trials with grating onset within 50 ms after microsaccades) and divided it by rate on no-microsaccade trials (i.e. trials with no microsaccades within <100 ms from grating onset). A value <1 indicates suppression.

Note that we only considered neurons with >5 spikes/s stimulus-evoked response (even on 11.11 cpd trials, which frequently had the lowest firing rates), thus avoiding “divide by zero” problems. Also, note that this modulation index isolates changes in visual sensitivity associated with microsaccadic suppression, regardless of how visual sensitivity itself might depend on spatial frequency without microsaccades. For example, visual responses in general are expected to be weaker for high spatial frequencies (Hafed and Chen 2016); however, our modulation index would normalize activity within a given spatial frequency in order to isolate any further suppression of visual sensitivity due to saccadic suppression.

In our analyses (including behavioral analyses), we also combined microsaccades towards or away from the grating because microsaccadic suppression is not direction-dependent in the post-movement interval that we focused on (Chen et al. 2015). We also confirmed this when analyzing the present data set (data not shown). We additionally considered a potential confound in our data. Specifically, because microsaccade trials were usually fewer than no-microsaccade trials for a given stimulus, it could be argued that the measured firing rates on microsaccade trials may have been biased relative to the no-microsaccade trials as a result of fewer trial repetitions. We thus checked whether the smaller numbers of trials for the microsaccade condition somehow explained our neural modulation results. For each neuron and spatial frequency, we subsampled the no-microsaccade trials, such that we kept only the same number of trials as in the corresponding microsaccade condition. We repeated such subsampling 10000 times with replacement, and we confirmed that all of our statistical results were not due to the microsaccade condition having fewer trials than the no-microsaccade condition (e.g. see Fig. 3). Finally, we combined neurons representing different eccentricities in our population analyses. We did so because we found that suppression is independent of eccentricity during the post-movement interval that we focused on (also see Chen et al. 2015).

To investigate the relevance of neural modulations to behavioral effects, we correlated behavioral patterns of saccadic suppression from the behavioral tests to neural modulations obtained from the recordings. For example, we related visual response strength to RT as a function of time of grating onset after microsaccades. We also did this for neural modulations of “first-spike latency”. This latency was defined as the time of visual burst onset after grating onset, and we investigated whether this time was affected by saccadic suppression. For neurons with no baseline activity (the majority), first-spike latency is trivially obtained as the first occurring spike after stimulus onset. For neurons with baseline activity, we used Poisson spike-train analysis to estimate visual burst onset (Legendy and Salcman 1985).

For all analyses with time courses, we used bin steps of 10 ms and bin widths of 50 ms (except for Fig. 2C, F with both bin steps and bin widths of 25 ms).

#### Local field potential (LFP) analyses

To analyze LFP’s, we sampled neurophysiological activity at 40 KHz. The signal was first filtered in hardware (0.7-6 KHz). We then removed 50, 100, and 150 Hz line noise using an IIR notch filter and then applied a zero-phase-lag lowpass filter (300 Hz cutoff). We finally down-sampled to 1 KHz. We analyzed filtered LFP traces like firing rates (Hafed and Chen 2016; Ikeda et al. 2015), and we classified electrode track locations as visual or visual-motor according to the neurons isolated from these tracks in the same sessions (Hafed and Chen 2016).

To obtain a measure of intrinsinc peri-microsaccadic modulation of LFP’s independent of visual stimulation, we took all microsaccades occurring in a pre-stimulus interval (20-100 ms before grating onset). We then aligned LFP traces on either microsaccade onset or end. To compare this data to a baseline, we took identically-long analysis intervals, again from pre-stimulus periods, but with no microsaccades occurring anywhere within these intervals.

To correlate LFP responses to behavioral dynamics of saccadic suppression (similar to what we did with firing rates), we measured peak transient LFP deflection as the minimum in the stimulus-evoked LFP trace 20-150 ms after grating onset. We created a “field potential index” by dividing this measurement on microsaccade trials by that on no-microsaccade trials. An index >1 indicates enhancement. For a control analysis, we computed the index after correcting for a microsaccade-related LFP level shift that may have happened due to intrinsic peri-microsaccadic modulation of the LFP independent of visual stimulation. We did this according to the following procedure. On microsaccade trials, we measured the average LFP value −25 to 25 ms from grating onset. We then subtracted the peak stimulus-evoked LFP deflection from this baseline measurement before dividing by the no-microsaccade trials. If an intrinsic peri-microsaccadic LFP modulation explained our results of LFP enhancement with increasing spatial frequency (see Results), then the baseline-shifted index should show no enhancement.

We also analyzed transient stimulus-evoked LFP deflection latency (analogous to first-spike latency). We found the first time at which the LFP was >2 s.d. away from baseline LFP (calculated as the mean LFP value −25 to 25 ms from grating onset), and there also had to be >5 ms of continuous >2 s.d. deviation from baseline. We did this separately for microsaccade and no-microsaccade trials, and we subtracted the measurements to obtain the influences of saccadic suppression on stimulus-evoked LFP deflection latency. If the LFP transient deflection occurs faster on microsaccade trials, then the subtraction gives a negative value.

## Results

### Selective Microsaccadic Suppression of Low Spatial Frequencies in Behavior

Isolation of perceptually-relevant saccadic suppression requires demonstrating a selective form of suppression in behavior, and subsequently asking which neurons reflect such selectivity. We thus first developed a sensitive behavioral measure demonstrating selective suppression, which was based on our earlier results (Hafed and Krauzlis 2010). We did so for microsaccades because they are mechanistically similar to larger saccades, while at the same time providing important experimental advantages (Materials and Methods). Monkeys fixated, and we initiated a computer process for real-time microsaccade detection (Chen and Hafed 2013). After such detection by a programmable delay, we presented a stationary vertical Gabor grating (80% contrast). The monkeys oriented towards the grating as fast as possible. Because SC *visual* bursts are strongly correlated with RT (Boehnke and Munoz 2008; Chen et al. 2015; Hafed and Chen 2016; Hafed et al. 2015; Hafed and Krauzlis 2010; Tian et al. 2016), we used RT changes in this task as a sensitive measure of microsaccadic influences on visual sensitivity (Hafed and Krauzlis 2010; Tian et al. 2016).

Similar to previously reported perceptual effects with large saccades (Burr et al. 1994) and microsaccades (Hass and Horwitz 2011), grating onset after microsaccades had a strong, yet selective, impact on behavior in our monkeys. Figure 1A shows example eye position (left) and velocity (right) traces from one monkey while we presented a 1.11 cpd grating. The black traces show trials without microsaccades <250 ms from grating onset, and the gray traces show trials with grating onset ~20-100 ms after microsaccades. There was a marked increase in RT during microsaccade trials (Fig. 1A). However, when we presented 4.44 cpd gratings, RT’s on microsaccade and no-microsaccade trials were similar (Fig. 1B; compare gray and black distributions). Thus, the microsaccadic suppressive effect (causing slower RT’s) was diminished for higher-frequency gratings. These sample-trial results demonstrate a microsaccadic correlate of selective perceptual suppression of low spatial frequencies by large saccades and microsaccades (Burr et al. 1982; Burr et al. 1994; Hass and Horwitz 2011; Volkmann et al. 1978).

**Figure 1.**
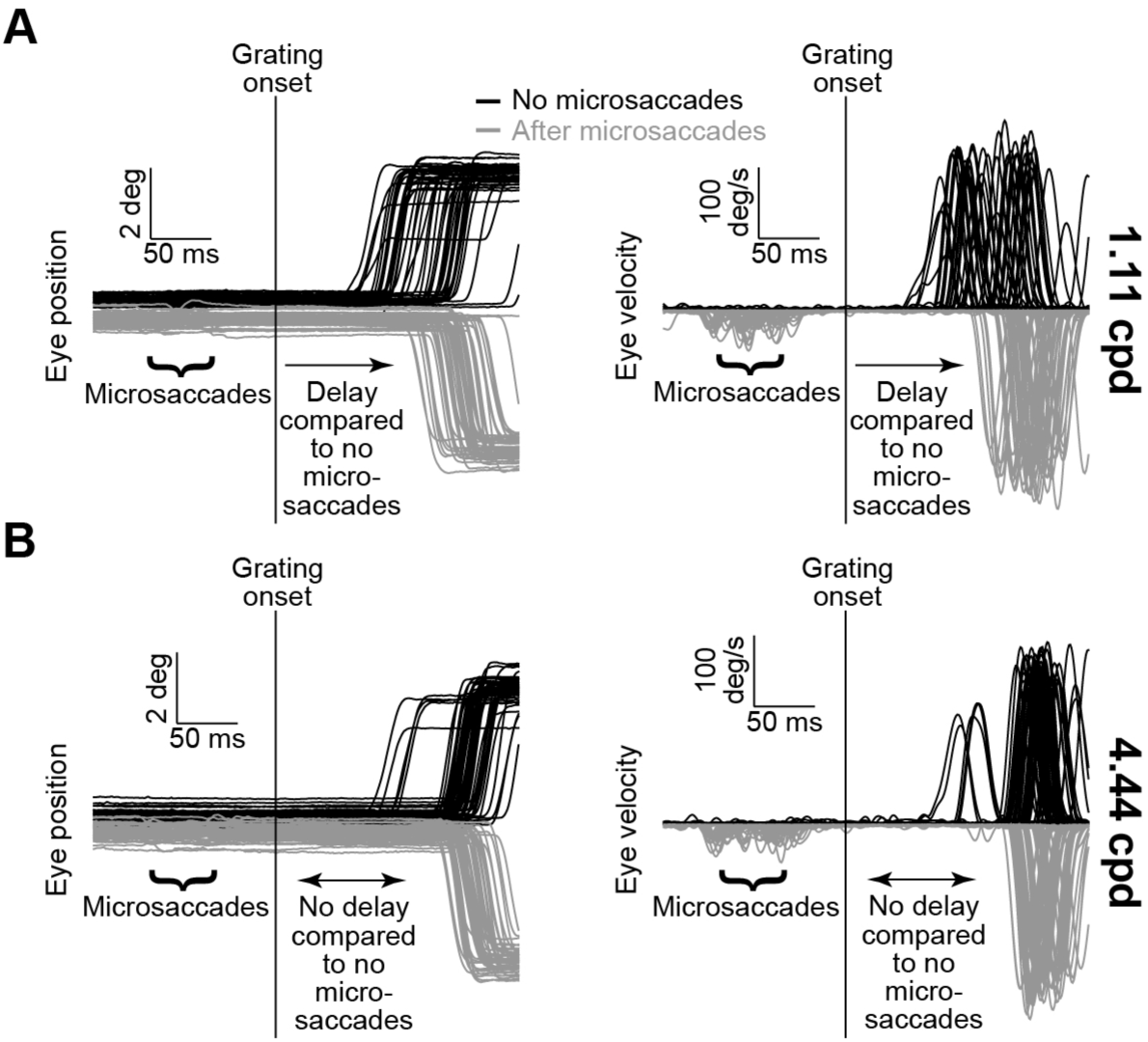
Behavioral measure of microsaccadic suppression across spatial frequencies. (**A**) Eye position (left) and radial eye velocity (right) traces from 100 sample trials from monkey N during a stimulus detection task. A 1.11 cpd grating appeared during fixation either with no nearby microsaccades (black, n=50 randomly selected trials) or ~20-100 ms after microsaccades (gray, n=50 randomly selected trials), and the monkey had to orient as fast as possible to the grating. Reaction time (RT) on the microsaccade trials was slower than on the no-microsaccade trials. Note that we flipped the gray position and velocity traces around the horizontal axis to facilitate comparison to the black traces, and we also displaced the initial fixation position in the position traces. The microsaccades are more visible in the velocity traces, because they constitute spikes of eye velocity. (**B**) Same analysis, but from 100 randomly selected trials having a higher spatial-frequency grating (4.44 cpd). RT’s in this case were more similar between the microsaccade and no-microsaccade trials, suggesting that the effect in **A** disappears with increasing spatial frequency.

Across sessions, both monkeys showed selective RT increases for low spatial frequencies (Fig. 2A, D). On no-microsaccade trials (black curves), RT increased with increasing spatial frequency, as expected from dynamics of the early visual system (Breitmeyer 1975) and SC (unpublished observations). This effect was statistically significant (p<0.01 for monkey N and p<0.01 for monkey P, 1-way ANOVA with spatial frequency as the main factor). However, with gratings appearing ~20-100 ms after microsaccades, the RT cost relative to no-microsaccade trials (i.e. the difference in RT between microsaccade and no-microsaccade trials) was strongest for the lowest spatial frequencies (Fig. 2B, E; p<0.01 for monkey N and p<0.01 for monkey P, 1-way ANOVA with spatial frequency as the main factor). This effect was not a ceiling effect on RT (because of the baseline increase with increasing spatial frequency). For example, at 4.44 cpd, RT on microsaccade and no-microsaccade trials was similar (Fig. 2A, D; magenta rectangles), but it got even slower for 11.11 cpd regardless of eye movements. Thus, the reduction in RT differences between microsaccade and no-microsaccade trials for high spatial frequencies (Fig. 2B, E) was indicative of a selective suppression of low spatial frequencies, and not necessarily a ceiling effect on RT.

**Figure 2.**
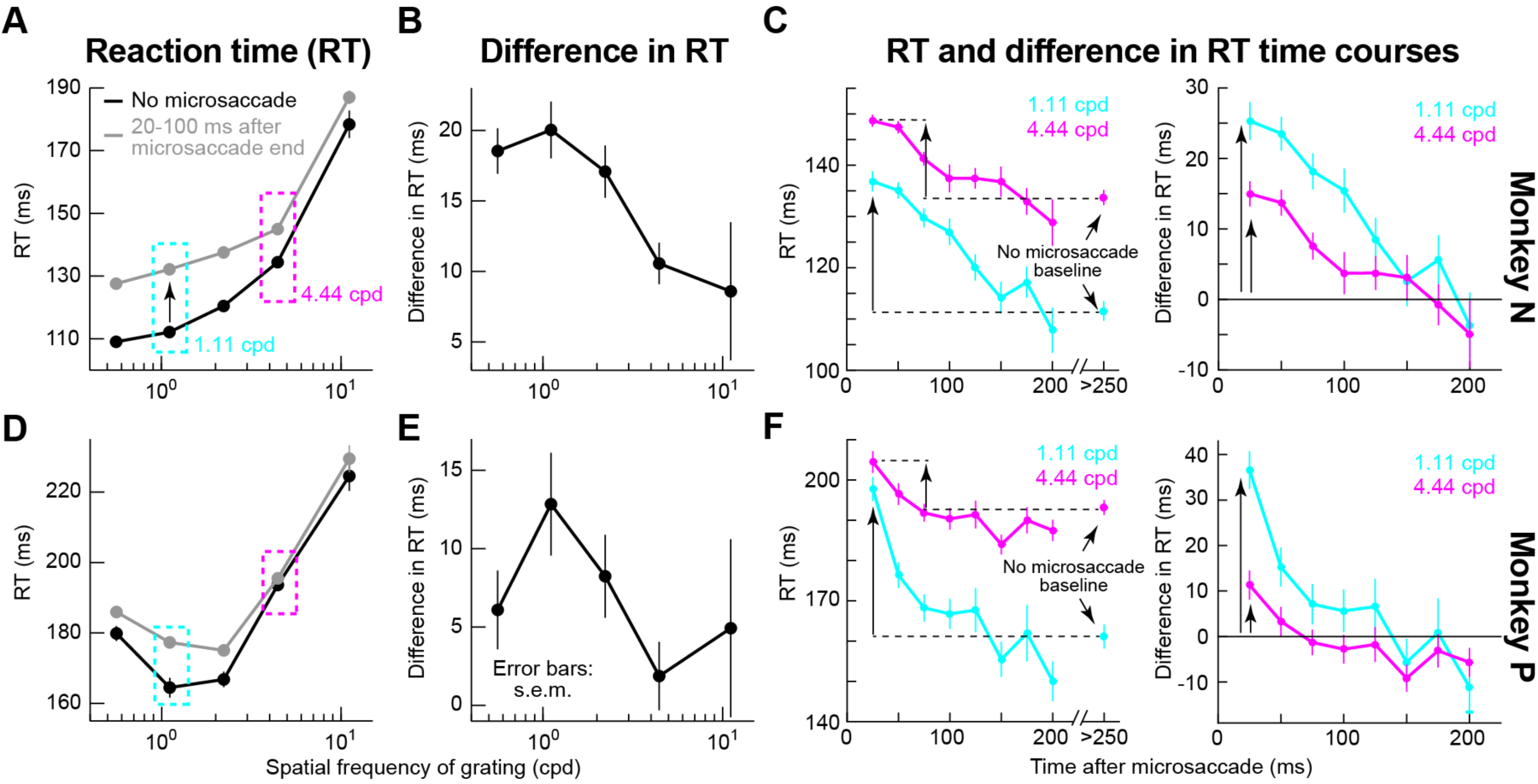
Spatial-frequency selective microsaccadic suppression in behavior. (**A**) RT as a function of spatial frequency. On no-microsaccade trials (black), RT increased with spatial frequency, consistent with dependence of visual response dynamics on spatial frequency (Breitmeyer 1975). If the same gratings appeared ~20-100 ms after microsaccades (gray), RT increased relative to no-microsaccade trials (a behavioral correlate of suppressed visual sensitivity), but more dramatically for low rather than high spatial frequencies (compare gray to black curves at different spatial frequencies). (**B**) Difference in RT between microsaccade and no-microsaccade trials (i.e. difference between gray and black curves in **A**), demonstrating the diminishing effects of microsaccades on RT behavioral costs with increasing spatial frequency. (**C**) Time courses of RT (left panel; like in **A**) or difference in RT (right panel; like in **B**) as a function of the time of grating onset after microsaccade end. The figure shows time courses from two sample spatial frequencies (complete time courses from all spatial frequencies, and for each animal individually, are also shown in Fig. 6). For the difference in RT time course, RT’s on trials with no microsaccades within <250 ms from grating onset were taken as the baseline. The initial RT cost caused by microsaccades was weaker for higher spatial frequency gratings (compare vertical arrows, consistent with **A**). Error bars, when visible, denote s.e.m. (**D-F**) Same analyses but for a second monkey. n=8153 trials for monkey N, and n=7117 for monkey P.

Our behavioral paradigm also provided rich information about saccadic suppression dynamics, which we could later use to identify a behaviorally-relevant SC neural modulation. For example, we evaluated microsaccadic suppression time courses across different spatial frequencies. Figure 2C, F illustrates this by plotting RT as a function of when a 1.11 or 4.44 cpd grating appeared after microsaccades. Microsaccadic suppression had a clear time course of RT costs for each spatial frequency, with both monkeys showing lower suppression for the higher spatial frequency immediately after microsaccades, and then a gradual return towards the baseline no-microsaccade performance for a given frequency.

Therefore, using a behavioral measure sensitive to SC visual response strength (Boehnke and Munoz 2008; Hafed and Chen 2016; Hafed et al. 2015; Hafed and Krauzlis 2010), we demonstrated a robust and selective pattern of microsaccadic suppression, which is directly analogous to perceptual suppression with large saccades (Burr et al. 1982; Burr et al. 1994; Hass and Horwitz 2011; Volkmann et al. 1978). We were now in a position to evaluate neural correlates of this suppression, and to specifically test a previously published hypothesis that perceptually-relevant suppression may emerge in purely visual SC neurons (Berman and Wurtz 2008; 2010; 2011; Isa and Hall 2009; Lee et al. 2007; Phongphanphanee et al. 2011; Wurtz 2008; Wurtz et al. 2011).

### Selective Suppression of Low Spatial Frequencies in Visual-Motor but not Visual SC Neurons

Using the same animals but in completely different experimental sessions not requiring any saccadic responses at all (Materials and Methods), we recorded the activity of purely visual SC neurons (24 neurons; located 680 +/- 95 s.e.m. μm below SC surface) or visual-motor neurons (66 neurons; 1159 +/- 66 s.e.m. μm below SC surface). Both types of neurons exhibit robust *visual* responses, but the question remains as to which would show perceptually-relevant suppression. We presented gratings similar to those used in Figs. 1-2 inside each neuron’s RF (Materials and Methods). However, the task was now a pure fixation task, and we only analyzed either no-microsaccade trials or trials in which the gratings appeared immediately *after* microsaccades (Materials and Methods).

Ensuring pure fixation was especially important to demonstrate behavioral relevance of our neural modulations. Specifically, one of our primary goals was to directly correlate neural dynamics to behavior in each animal (as will be presented later). Showing that a specific SC cell class is highly correlated with behavior compared to another cell class, *even* when the correlations are made across completely independent sessions and tasks, would demonstrate the behavioral relevance of the cell class. Moreover, demonstrating that neural suppression dynamics appear on *visual* responses, even in the complete absence of an overt response, shows that it is *sensory* responses that matter during saccadic suppression. Finally, ensuring fixation avoided saccade preparation influences on visual sensitivity (Li and Basso 2008).

Visual-motor SC neurons showed the strongest saccadic suppression, and in a spatial-frequency selective manner. Figure 3A shows the activity of two sample pure visual neurons (one per row) during presentations of different spatial frequencies (across columns), and Fig. 3B shows the activity of two sample visual-motor neurons (in the same format). In each panel, saturated colors show activity with no microsaccades <100 ms from grating onset, and unsaturated colors show activity when the same grating was presented within 50 ms after microsaccades. In no-microsaccade trials, all neurons showed expected visual bursts, but burst strength varied with spatial frequency (Fig. 3, saturated colors). This is suggestive of spatial-frequency tuning (Hafed and Chen 2016), but our purpose here was to investigate suppression relative to no-microsaccade responses; thus, we scaled the y-axis in each panel such that across panels, no-microsaccade curves visually appeared to be roughly equal in height. Using such scaling, visual-burst suppression (unsaturated colors) was rendered clearer (and quantitatively, we always measured suppression relative to the no-microsaccade responses *within* each given spatial frequency independently and not across spatial frequencies; Materials and Methods). Importantly, there were differences in suppression patterns between visual and visual-motor neurons. For the visual neurons (Fig. 3A), suppression was mild and relatively inconsistent across spatial frequencies; for the visual-motor neurons (Fig. 3B), there was strong suppression for the lowest spatial frequency (Neuron #3: ~32%; Neuron #4: ~38%; p<0.01 for each neuron, Mann-Whitney U-test), and there was also a systematic reduction in suppression strength with increasing frequency (by 4.44 and 11.11 cpd, there was practically no suppression left at all; p=0.49 for 4.44 cpd and p=0.41 for 11.11 cpd in Neuron #3, and p=0.15 for 4.44 cpd and p=0.99 for 11.11 cpd in Neuron #4, Mann-Whitney U-test). Importantly, the eye movement associated with suppression in all panels had ended before grating onset. Thus, the suppression cannot be attributed to blurring of the gratings by eye movements. Moreover, the suppression cannot also be due to a reduced number of trial repetitions in the microsaccade condition: the shaded curves in Fig. 3 show no-microsaccade firing rates after we subsampled the no-microsaccade condition to match trials with the microsaccade condition (Materials and Methods); the firing rates were statistically indistinguishable from our original full set of no-microsaccade trials.

**Figure 3.**
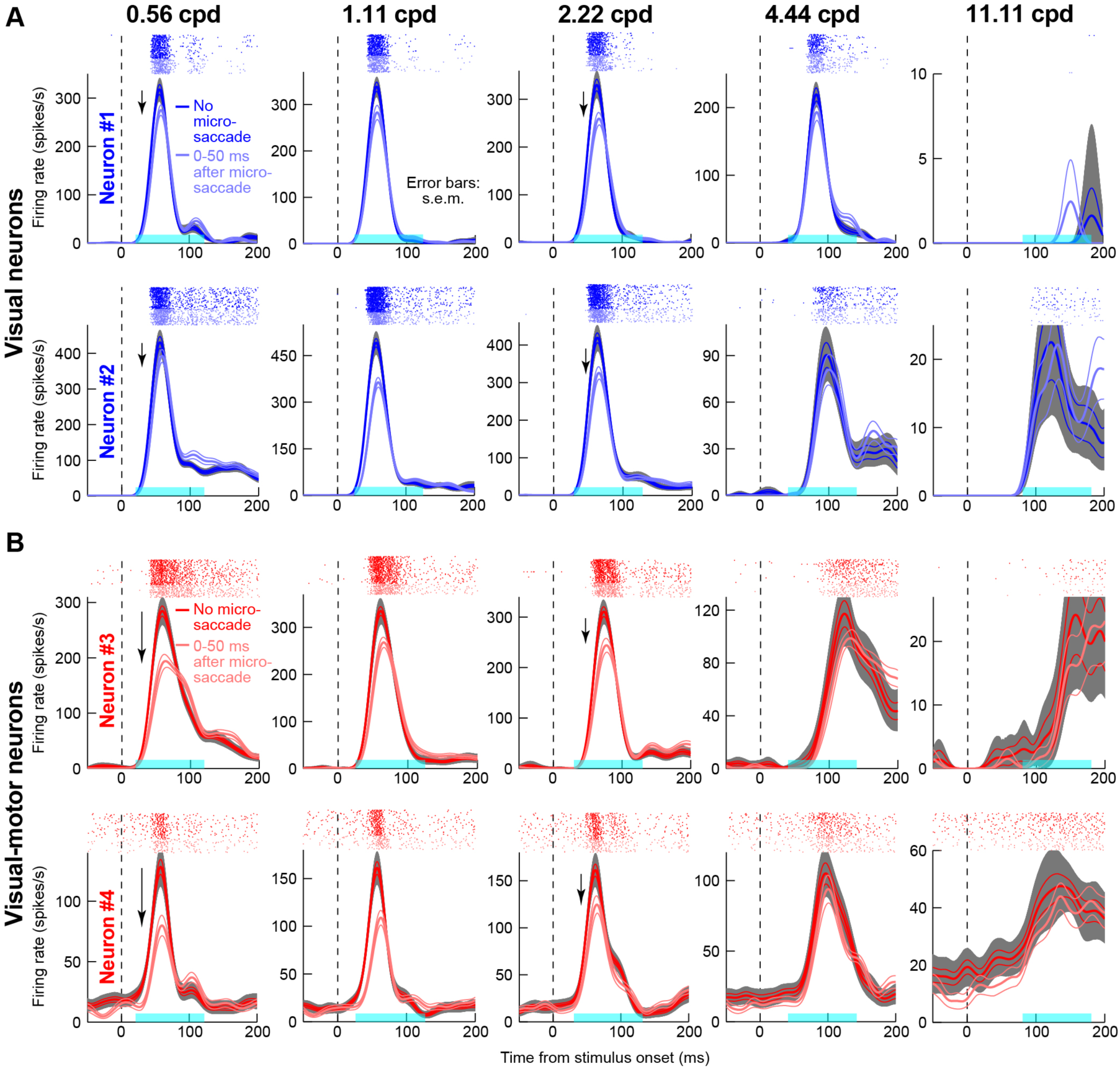
Spatial-frequency selective microsaccadic suppression of visual-motor SC neurons. (**A**) Neural activity as a function of time after grating onset for two sample purely visual SC neurons (one per row). Each panel in a row shows activity after presentation of a specific spatial frequency (indicated above each panel). Rasters above each firing rate curve show individual action potentials emitted by the neuron across individual trials. We divided trials into ones in which there was no microsaccade within <100 ms from grating onset (saturated blue; n>=38 trials per spatial frequency in these sample neurons) and ones in which the grating appeared immediately after microsaccades (unsaturated blue; n>=30 trials per spatial frequency). The y-axis was scaled in each panel such that the no-microsaccade firing rates visually appeared to have approximately similar heights across panels, allowing easier comparison of suppression effects. Both neurons showed moderate microsaccadic suppression, with no clear pattern across spatial frequencies. (**B**) Same format as **A**, but for two sample visual-motor neurons. The neurons showed stronger suppression at the lowest spatial frequency, and the suppression gradually decreased in strength with increasing spatial frequency (like in behavior); by 4.44 and 11.11 cpd, there was practically no suppression left at all. For these neurons, n>=28 trials per spatial frequency for no microsaccade trials (saturated red), and n>=22 trials per spatial frequency trials for microsaccade trials (unsaturated red). Error bars denote s.e.m. In all panels, the gray shaded regions indicate 95% confidence intervals for the no-microsaccade trials when they were subsampled (and bootstrapped) to match the numbers of trials in the microsaccade condition (Materials and Methods). Also, the cyan intervals on x-axes indicate the averaging intervals for measuring mean firing rate (Materials and Methods).

Across neurons, there was selective suppression of visual sensitivity as a function of spatial frequency, but only in visual-motor neurons. Figure 4 summarizes these findings by plotting a suppression index (Materials and Methods) as a function of spatial frequency. Visual bursts were suppressed in both visual and visual-motor neurons (suppression index <1). However, the suppression was not spatial-frequency selective, and it was weaker, in visual neurons; in visual-motor neurons, there was strong suppression only for the lowest spatial frequencies, and the effect gradually dissipated away with increasing frequency. We confirmed these observations statistically for both peak and mean visual response (i.e. both Fig. 4A and Fig. 4B). One-way ANOVA’s revealed a main effect of spatial frequency in visual-motor (p<0.01) but not visual (p=0.29) neurons. Also, the average suppression in visual neurons was 11% across spatial frequencies (p=0.1, Mann-Whitney U-test), and it was 22% in visual-motor neurons (p<0.01, Mann-Whitney U-test). A difference between visual and visual-motor neurons also appeared in suppression temporal dynamics (Fig. 5). Thus, there are differences in saccadic suppression strength between visual and visual-motor SC neurons, and visual-motor neuron suppression selectivity appears more similar to behavioral effects, both in our own experiments (Figs. 1-2) as well as in the literature of human perceptual effects (Burr et al. 1982; Burr et al. 1994; Volkmann et al. 1978).

**Figure 4.**
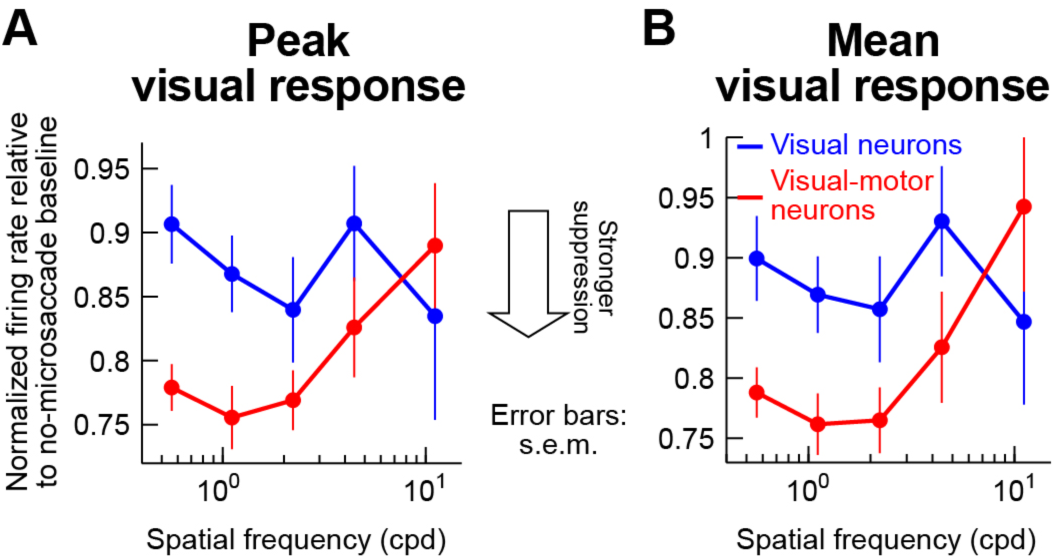
Spatial-frequency dependent microsaccadic suppression of visual bursts in visual-motor but not visual SC neurons. (**A**) We measured peak stimulus-evoked visual burst after grating onset (e.g. from traces like those in Fig. 3) and plotted it as a function of grating spatial frequency. We grouped neurons as purely visual (blue) or visual-motor (red). Visual neurons showed only ~10% suppression, and there was no consistent spatial-frequency dependence of this suppression. Visual-motor neurons showed ~25% suppression in the low spatial frequencies, and this effect gradually decreased with increasing spatial frequency (as in behavior). Error bars denote s.e.m. Note that the error bars for the highest spatial frequency were larger than other frequencies because some neurons completely stopped responding at 11.11 cpd, which reduced population size in this spatial frequency (Materials and Methods). (**B**) We repeated the same analysis but now measuring mean stimulus-evoked response in a certain window after grating onset (cyan intervals indicated in Fig. 3; Materials and Methods). Similar observations to those in **A** were made. n=66 visual-motor neurons, and n=24 visual neurons.

**Figure 5.**
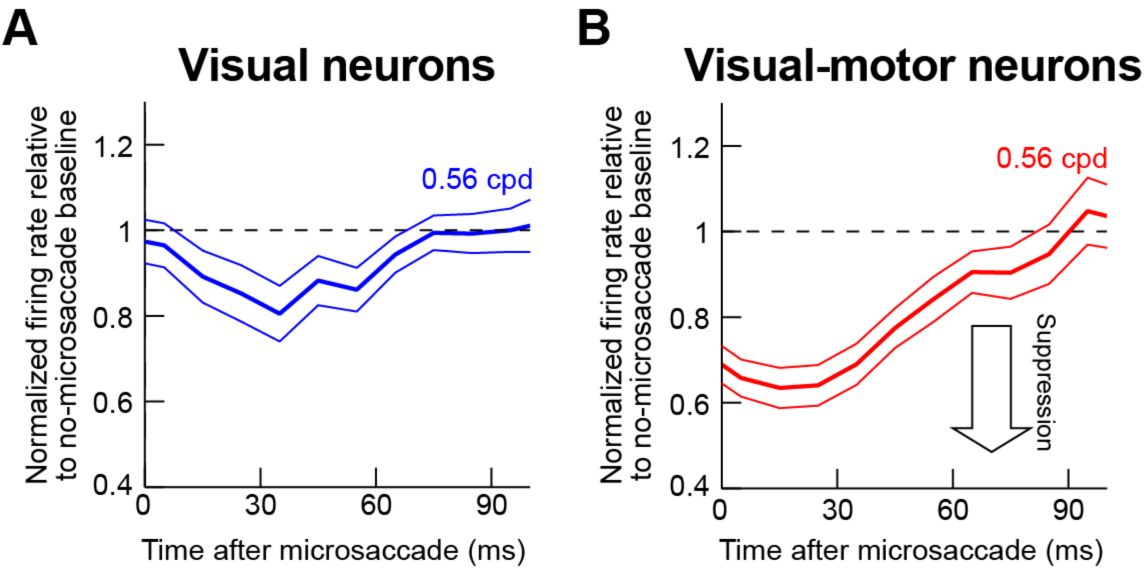
Time courses of microsaccadic suppression in visual (**A**) and visual-motor (**B**) neurons for a sample spatial frequency. We performed an analysis similar to that described in (Chen et al. 2015) but aligning on microsaccade end. For each time window after microsaccade end in which a grating appeared (x-axis; 50-ms bins in 10-ms steps), we measured mean firing rate evoked by grating onset (Materials and Methods), and we normalized it by mean firing rate on no-microsaccade trials. Visual-motor neurons showed stronger suppression than visual neurons (compare y-axis in both panels; error bars denote 95% confidence intervals), and both neuron types experienced recovery with increasing time after microsaccades (consistent with behavioral effects). Note that the time course of visual-motor neuron suppression is similar to the time course of behavioral effects (e.g. Fig. 2C, F) and also similar to the time course of saccadic suppression in the earlier literature (e.g. Ibbotson and Krekelberg 2011; Hafed and Krauzlis 2010; Diamond et al. 2000). Figure 6 shows individual monkey time courses, other spatial frequencies, as well as relationships between neural time courses and the respective monkey’s behavioral performance dynamics.

### Better Correlation Between Visual-Motor Neuron Dynamics and Behavior than Between Visual Neuron Dynamics and Behavior

To further explore the apparent similarity between visual-motor neuron suppression patterns (Fig. 4) and behavior (Fig. 2), we used the dynamics of our recorded population as a proxy for how the SC might be engaged in our behavioral task of Figs. 1-2. We plotted the time course of behavioral suppression (similar to Fig. 2C, F) for each spatial frequency and each monkey individually (Fig. 6A, E), and we also plotted the neural time course of visual-motor neuron suppression, again for each monkey individually (Fig. 6B, F; an example time course for purely visual neurons can also be seen in Fig. 5A). For this comparative analysis, we used the same binning windows in both behavioral and neural data (50-ms bin widths in steps of 10 ms starting at 0 ms after microsaccade end), and we next correlated the two time courses: we plotted all samples of the behavioral time course against all samples of the neural time course irrespective of spatial frequency or time after microsaccades (Fig. 6C, G). There was high correlation between visual burst strength in SC visual-motor neurons and the behavioral effect of microsaccadic suppression: whenever visual bursts were weaker, RT costs increased, and vice versa, regardless of spatial frequency or time after microsaccades. This high correlation is particularly remarkable given that the behavioral and neural data were collected in completely different sessions and with different behavioral tasks.

**Figure 6.**
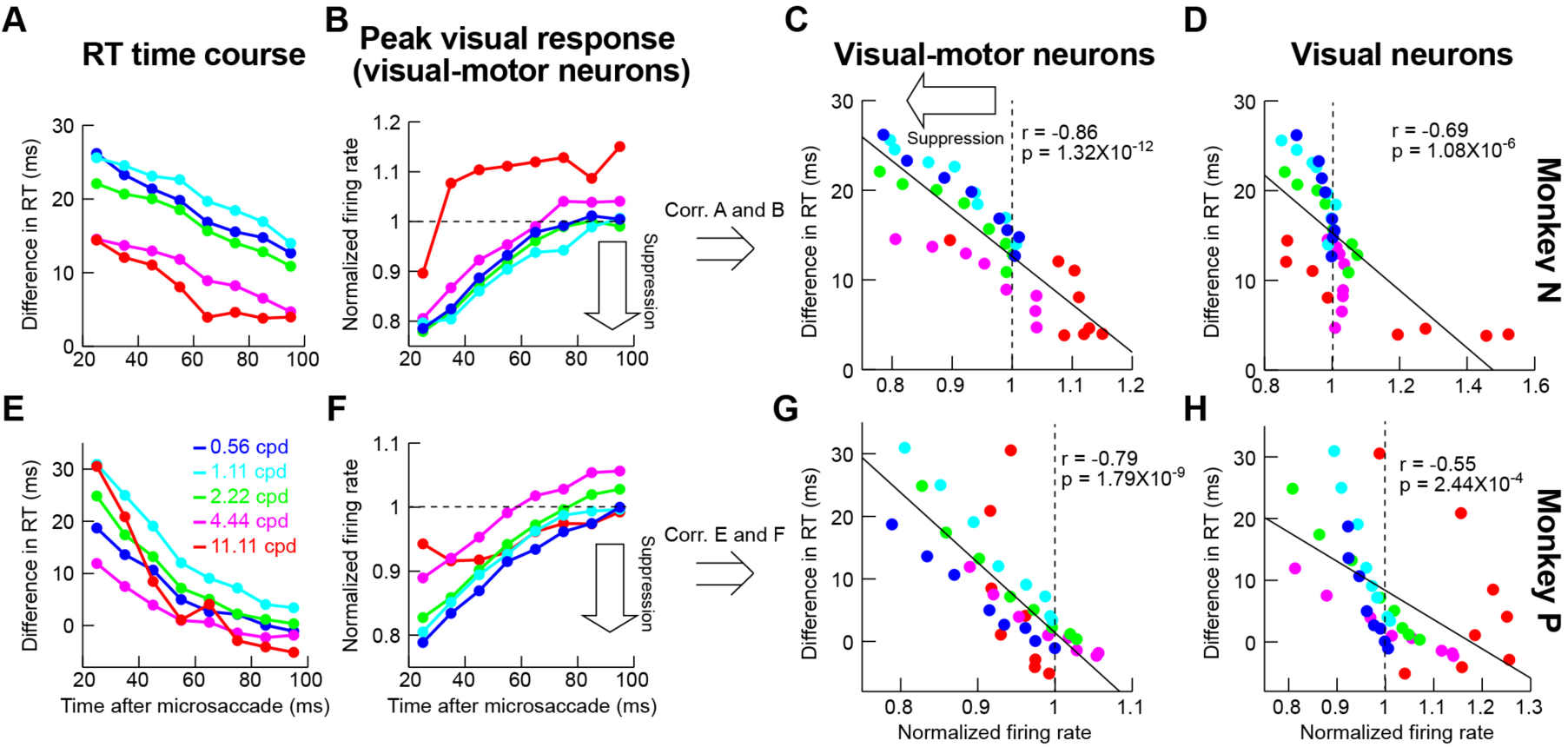
Correlating behavioral microsaccadic suppression with neural microsaccadic suppression on completely different experimental sessions. (**A**) Time course of difference in RT from baseline (e.g. Fig. 2C, F; Materials and Methods) as a function of time of grating onset after microsaccade end in our behavioral experiments (monkey N). Different curves show different spatial frequencies. Immediately after microsaccades, there was a strong cost in RT for low spatial frequencies, and a more moderate cost for high spatial frequencies. In all spatial frequencies, the RT cost associated with microsaccadic suppression slowly dissipated in time. (**B**) Similar analysis but for the peak visual response in our neural experiments, on completely different sessions from the behavioral data, and only for visual-motor neurons. (**C**) Correlation between the data points in **A** and those in **B**, regardless of time or spatial frequency. There was strong correlation between visual burst strength and RT cost, even on completely different experimental sessions, suggesting that visual-motor neurons are modulated during microsaccadic suppression in a perceptually-relevant manner. (**D**) This was not the case for purely visual neurons. Here, we correlated the behavioral points in **A** with similar points but for visual neuron time courses (e.g. Fig. 5A). The correlation with behavior was worse than in visual-motor neurons. (**E-H**) Similar observations for a second monkey. n=24 visual-motor neurons for monkey N, and n=42 visual-motor neurons for monkey P; n=15 visual neurons for monkey N, and n=9 visual neurons for monkey P.

The highest correlation between neural patterns and behavior was observed only when we used peak visual response of visual-motor SC neurons as the behavioral predictor (Fig. 6C, G). When we correlated behavioral time courses with peak visual response of purely visual neurons, the correlations were significantly worse (Fig. 6D, H; p=0.02 for monkey N and p=0.02 for monkey P, Steiger’s Z-test). We also tried to correlate behavior to mean visual response after stimulus onset (within a certain time window; Materials and Methods) or first-spike latency (whether of visual or visual-motor neurons), but peak visual response of visual-motor neurons always provided the best predictor. Specifically, even though there was a modest change in first-spike latency when visual bursts were suppressed (e.g. see Fig. 3), the effect was not as consistent as the effect of visual response magnitude, and it meant that neural response latency was not as good a correlate of behavioral effects as peak visual response.

The results of Fig. 6 suggest that saccadic suppression in visual-motor neurons is more in line with behavioral effects than for purely visual neurons. This is inconsistent with a hypothesized role of a feedback pathway from visual-motor to visual neurons in perceptually-relevant saccadic suppression in the SC (Isa and Hall 2009; Lee et al. 2007; Phongphanphanee et al. 2011). However, one possible confound could be in the distribution of preferred spatial frequencies in visual-motor neurons. For example, if only the preferred spatial frequency of a neuron experiences the strongest suppression, and if visual-motor neurons only had low preferred spatial frequencies, then the selective suppression of Fig. 4 would emerge. However, we found no clear differences in patterns of preferred spatial frequencies between visual and visual-motor neurons. Moreover, we explicitly analyzed suppression profiles of visual-motor neurons as a function of the neurons’ preferred spatial frequencies. For each spatial frequency, we took only neurons preferring this spatial frequency, and we checked how these neurons were suppressed.

Figure 7A-D shows the results of this analysis. There was indeed a tendency for the preferred spatial frequency of a neuron to experience the strongest suppression relative to other frequencies (e.g. black arrows). However, this strongest suppression still became progressively weaker and weaker with increasing spatial frequency (e.g. compare Fig. 7A, B to Fig. 7C, D). This is further demonstrated by Fig. 7E, in which we took the maximal suppression frequency from each of the panels in Fig. 7A-D and plotted them with an indication of the behavioral microsaccadic suppression profile (obtained as the inverse of RT profiles from Fig. 2B, E, with arbitrary y-axis scaling). Importantly, we again made sure that the neural suppression data in this figure were analyzed in an identical manner to behavioral analyses (i.e. we considered the same interval of stimulus onsets happening 20-100 ms after microsaccade end as in the behavioral analyses). As can be seen, there was a clear match between neural and behavioral effects in both animals (the correlation between neural suppression and behavioral suppression in this figure was 0.99 for monkey N and 0.89 for monkey P). Thus, the selective suppression of Figs. 3-6 was not an artifact of potential biased spatial-frequency tuning properties of only visual-motor neurons.

**Figure 7.**
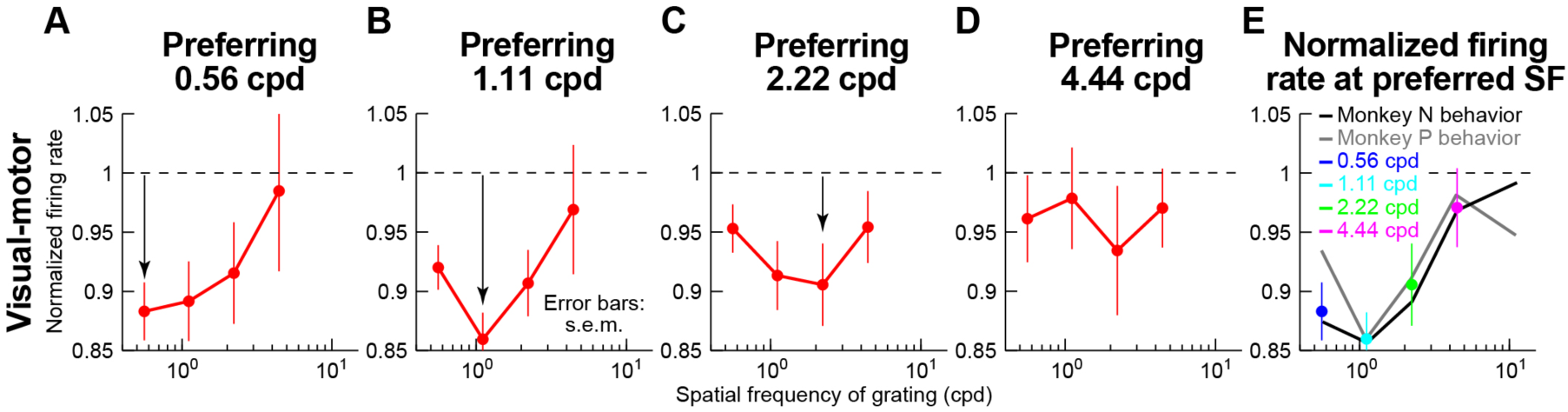
Selective low-frequency suppression in visual-motor neurons independent of preferred spatial frequency. (**A-D**) In each panel, we selected only neurons preferring a single spatial frequency on no-microsaccade trials (Materials and Methods). We then repeated the analysis of Fig. 4A. The preferred spatial frequency tended to experience the strongest suppression compared to other spatial frequencies (black arrows). However, the strength of the suppression even for the preferred spatial frequency consistently decreased with increasing spatial frequency (compare the arrows in the individual panels). Note that we did not have neurons preferring 11.11 cpd in this analysis, and we thus do not show this spatial frequency in this figure. (**E**) We collected the maximally suppressed spatial frequency from each panel in **A**-**D** (legend), and we plotted them together. The black and gray lines are a copy of the behavioral RT microsaccadic suppression curves of Fig. 2B, E, but inverted (and with arbitrary y-axis scaling) to match the neural suppression curves. As can be seen, even if the preferred spatial frequency of neurons always experienced maximal suppression, this maximal suppression was still decreased with increasing spatial frequency. Thus, the spatial-frequency selectivity of visual-motor neural suppression was still correlated with behavior. Error bars denote s.e.m. n=26, 19, 8, and 5 neurons in each of **A**, **B**, **C**, and **D**.

Taken together, our results so far suggest that perceptually-relevant SC saccadic suppression (i.e. selective for spatial frequency as in perception) is localized in the visual-motor neurons, with visual neurons only showing modest and non-selective modulations.

### Influence of a Putative Microsaccadic Source Signal on Local SC Population Activity During Suppression

To demonstrate that there may indeed be a saccadic source signal associated with suppressed SC visual bursts (i.e. putative corollary discharge), we analyzed local field potentials (LFP’s) around our electrodes (Materials and Methods). Stimulus onset in no-microsaccade trials caused a negative-going “stimulus-evoked” LFP deflection for both visual and visual-motor electrode tracks (Hafed and Chen 2016; Ikeda et al. 2015). For example, Fig. 8 shows LFP traces (in a format similar to Fig. 3) as a function of spatial frequency for an example superficial track (i.e. among visual neurons; Fig. 8A) and an example deeper track (among visual-motor neurons; Fig. 8B). Remarkably, on microsaccade trials (unsaturated colors), stimulus-evoked LFP response was not suppressed. In fact, for the visual-motor electrode track (Fig. 8B), LFP response was enhanced, and more so with increasing spatial frequency (Fig. 9A; p<0.01, 1-way ANOVA on the modulation index with spatial frequency as the main factor). Given that LFP’s reflect not just local population spiking activity, but also putative synaptic inputs, these results suggest the existence of a possible microsaccade-related input modulating visual bursts, and this effect was again stronger in visual-motor than visual electrode tracks (Fig. 9A). However, it is important to emphasize here that this signal was not a direct microsaccade command because none of our neurons at all electrode locations in this study exhibited microsaccade-related movement bursts (Materials and Methods).

**Figure 8.**
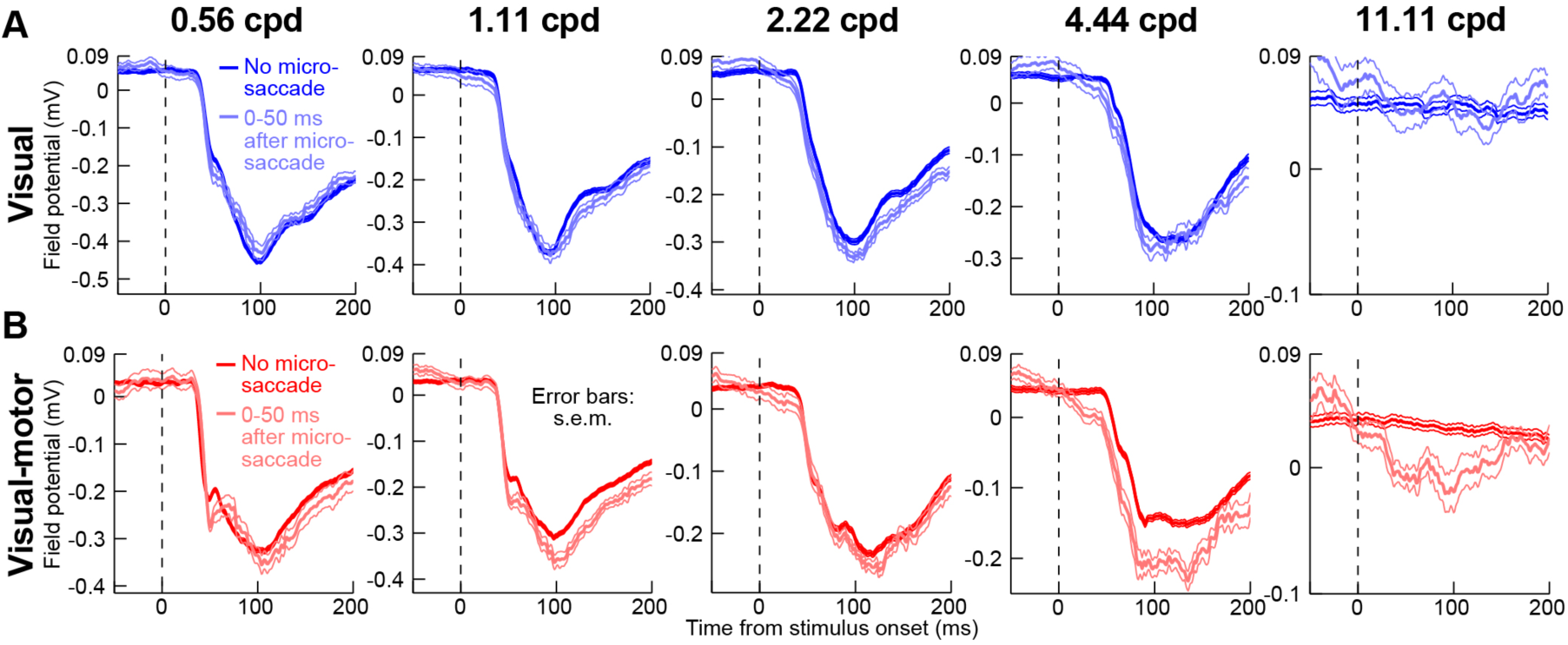
Local field potential modulations during microsaccadic suppression. This figure is formatted similarly to Fig. 3, except that we now plot LFP modulations around a sample electrode track near visual (**A**) or visual-motor (**B**) neurons. There was *no* evidence of a reduced LFP evoked response for trials with grating onset after microsaccades (faint colors). If anything, the peak evoked response, and the latency to evoked response were stronger and shorter, respectively (see Fig. 9). This effect was not explained by an intrinsic peri-microsaccadic modulation of LFP (see Figs. 9A, 10), but it is consistent with an additional movement-related modulatory signal associated with saccade execution that influences stimulus-evoked spiking activity. Error bars denote s.e.m. For the visual track (**A**), n>=113 trials per spatial frequency on no-microsaccade trials (saturated blue), and n>=25 trials per spatial frequency on microsaccade trials (unsaturated blue). For the visual-motor track (**B**), n>=140 trials per spatial frequency on no-microsaccade trials (saturated red), and n>=12 trials on microsaccade trials (unsaturated red).

**Figure 9.**
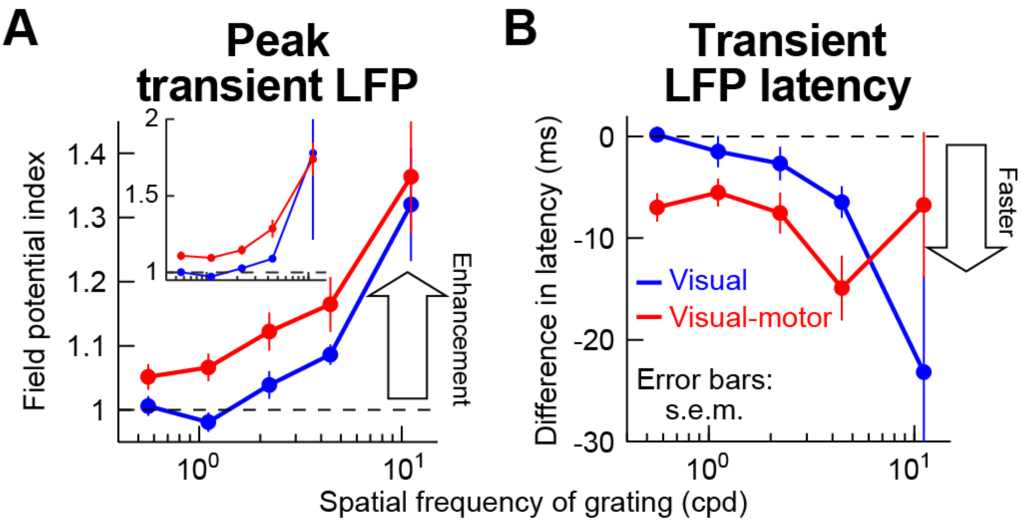
Lack of microsaccadic suppression in LFP stimulus-evoked responses. (**A**) We performed an analysis similar to that in Fig. 4 but on LFP’s. We measured peak LFP response with and without microsaccades, and we then obtained a modulation index (Materials and Methods). The inset shows the modulation index from raw measurements, and the main panel shows the same analysis but after subtracting a baseline shift from the microsaccade trials. Specifically, Fig. 10 suggests that there is a negativity in LFP’s after microsaccades, and stimulus onset in the microsaccade trials came after a previous microsaccade. Thus, we measured the peak LFP stimulus-evoked response on microsaccade trials as the difference between the raw LFP stimulus-evoked negativity minus the baseline LFP value that was present at the time of grating onset (Materials and Methods). In both the inset and the main panel, there was no suppression in the stimulus-evoked LFP response, contrary to firing rate results (Fig. 4). Rather, there was response enhancement, which progressively increased with increasing spatial frequency, and this happened for both visual and visual-motor electrode track locations (p<0.01 for either baseline-corrected or raw measurements and for each of visual-only or visual-motor electrode tracks; 1-way ANOVA with spatial frequency as the main factor). (**B**) Similar analyses but measuring the latency to LFP stimulus-evoked response, which decreased on microsaccade trials (y-axis values <0 ms; p<0.01 for visual electrode tracks and p=0.03 for visual-motor electrode tracks; 1-way ANOVA with spatial frequency as the main factor). Thus, when a stimulus appeared immediately after a previous microsaccade, the stimulus-evoked LFP response started earlier than without a microsaccade. Error bars denote s.e.m.

Our interpretation of a modulatory movement-related input mediating firing rate suppression effects is consistent with the enhanced LFP effect seen in Figs. 8 and 9A for high spatial frequencies. These frequencies evoke the weakest visual activity (Figs. 3, 8; saturated colors), meaning that the influence of a saccadic source signal for suppression (which is dependent on the movement and not the stimulus) should become increasingly more obvious in the LFP with increasing spatial frequency (Fig. 9A). Thus, combined with earlier firing rate results, our LFP analyses reveal that visual-motor SC neurons may be closely associated with a movement-related source for perceptually-relevant saccadic suppression.

One possible confound with the above result is that microsaccades (even though they ended before stimulus onset) might cause long-lasting LFP modulations, which would be superimposed on a stimulus-evoked LFP deflection in Fig. 8. In other words, the evoked response could potentially still be suppressed, but it could be level-shifted because it rides on a microsaccade-induced LFP modulation. Indeed, during simple fixation without any other visual stimuli, both visual and visual-motor SC electrode locations exhibited prolonged microsaccade-related LFP modulations, involving a subtle negativity after microsaccade end (Fig. 10). It is intriguing that this effect happens even in extra-foveal SC (i.e. with no microsaccade-related bursting neurons), and in purely visual layers as well, because it suggests that the modulation is not a direct motor command. Instead, it is a modulatory effect that is far-reaching across the SC, potentially explaining previously observed peri-microsaccadic modulations in neural activity and behavior (Chen et al. 2015; Hafed 2013; Hafed et al. 2015; Tian et al. 2016). The modulation is also similar to saccade-related LFP modulations in human SC (Liu et al. 2009). In any case, what this modulation (Fig. 10) means for Fig. 8 is that an LFP negativity following microsaccades might alter the baseline on top of which the stimulus-evoked deflection rides on.

**Figure 10.**
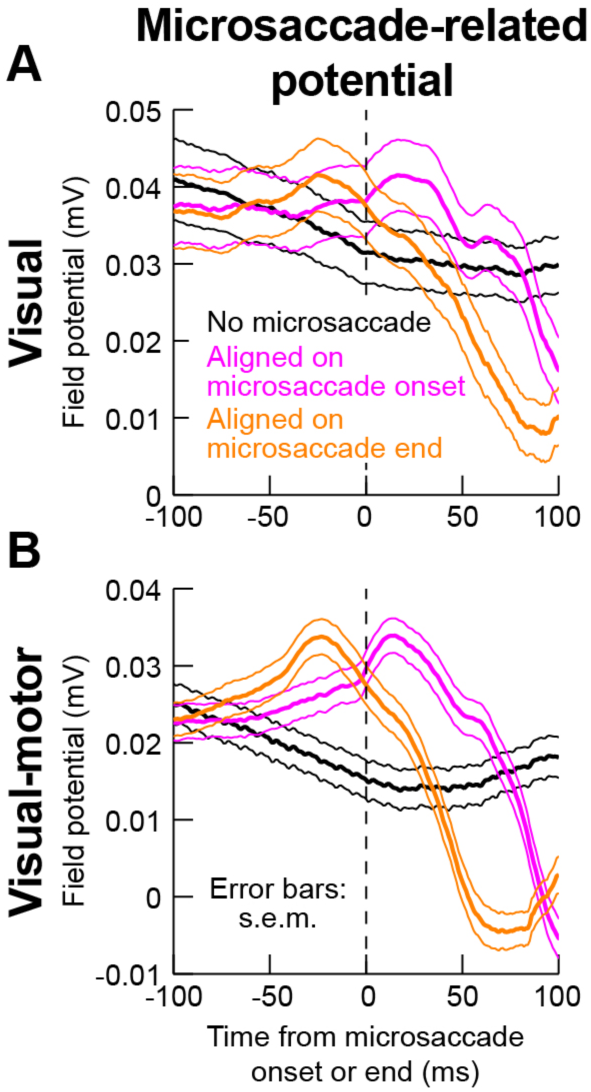
Microsaccade-related local field potential modulations in the absence of an RF stimulus. We aligned LFP activity to either microsaccade onset (magenta) or microsaccade end (orange) during a baseline fixation interval with no RF stimulus at all (Materials and Methods). The black curves show LFP activity during equally-long control intervals, again with no RF stimulus, but also with no microsaccade occurrence. Even though there was no microsaccade-related spiking at all the sites investigated in this study, microsaccades caused systematic modulations in both visual (**A**) and visual-motor (B) electrode locations in the SC, even though our electrodes were primarily placed in extra-foveal SC representations far from the movement endpoints. Thus, these LFP modulations, similar to previously reported saccade-related LFP modulations (Liu et al. 2009), reflect a potential microsaccade-related modulatory signal that can mediate microsaccadic suppression of firing rates in extra-foveal SC neurons. Also, note how the effect on visual-motor layers (**B**) is more systematic and robust than in visual layers (**A**). This is further evidence of a putative extra-retinal signal in the SC visual-motor layers that might mediate saccadic suppression (and explain Fig. 6), and it also makes it unlikely that the LFP modulations in this figure are due to ocular muscle artifacts. Error bars denote s.e.m. n=66 electrode tracks for (**B**), and n=24 electrode tracks for (**A**).

However, we found that the low amplitude of peri-microsaccadic LFP modulation (Fig. 10) was not sufficient to explain the lack of LFP suppression in stimulus-evoked LFP’s (Fig. 8). Specifically, we corrected for a baseline shift at grating onset (Materials and Methods), and we still found no suppression in the strength of the stimulus-evoked LFP response (Fig. 9A). Thus, in Figs. 8-10, we believe that we have uncovered evidence for a putative microsaccade-related modulatory input at the time of visual burst suppression in both SC visual and visual-motor neurons. Moreover, this input shows differential modulation between superficial and intermediate electrode tracks (Fig. 9A), consistent with our firing rate results.

Enhanced stimulus-evoked LFP response amplitudes (Fig. 9A) were also accompanied by slightly faster LFP responses (Fig. 9B), again consistent with a movement-related source modulating neural firing rates at the time visual burst occurrence (because the movement happened before stimulus onset). It is also interesting to note that, like firing rate time courses, time courses of stimulus-evoked LFP modulations for stimuli appearing after microsaccades were also correlated with behavioral microsaccadic suppression dynamics (as in Fig. 6). In the LFP’s, the best behavioral predictor was the latency of stimulus-evoked LFP deflection, and visual-motor electrode tracks again showed higher correlation values with behavior than visual electrode tracks. These results are shown in Fig. 11, which is formatted similarly to Fig. 6, except that the neural data in this figure was now based on LFP measurements instead of firing rates.

**Figure 11.**
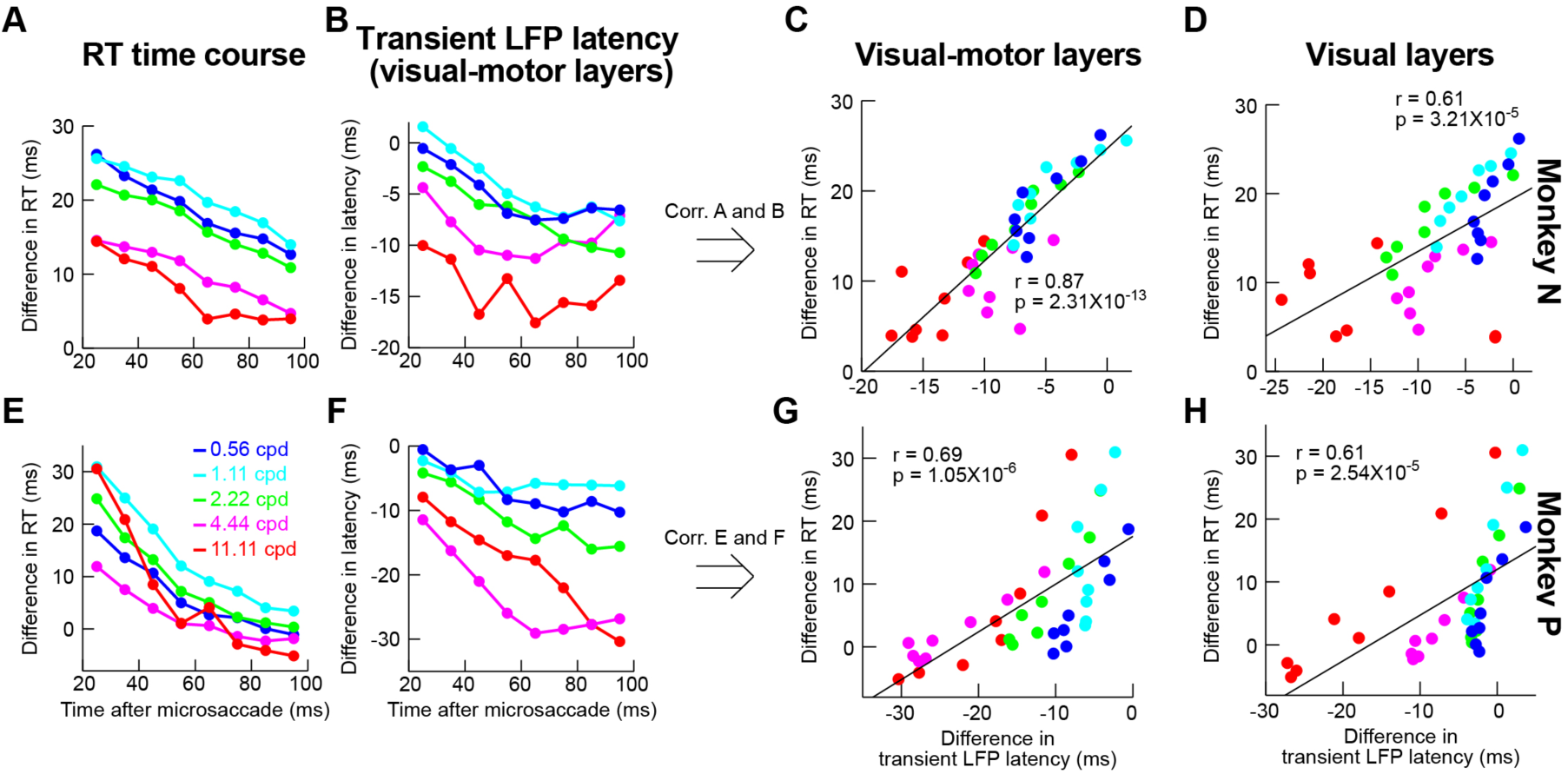
Correlation between LFP modulation parameters and behavioral effects of suppression. This figure is formatted similarly to Fig. 6, except that here we plotted LFP time courses instead of firing rate time courses. Specifically, in **B** and **F**, we plotted the time course of LFP stimulus-evoked response latency (e.g. Fig. 9B) as a function of spatial frequency and time after microsaccades. The correlation between this latency in visual-motor layers and behavior was better (**C, G**) than in visual layers (**D, H**). Thus, it is again the visual-motor layers that are better predictors of behavior, like in Fig. 6, although firing rates (Fig. 6) showed higher correlations to behavior in general. Note that we also measured correlations between behavior and LFP stimulus-evoked response strength rather latency (data not shown), but the LFP response latency always showed the better correlations with behavior.

Our results combined demonstrate that visual-motor neurons are more in line with selective perceptual effects of saccadic suppression, both in humans (Burr et al. 1982; Burr et al. 1994; Volkmann et al. 1978) and monkeys (Fig. 2), than purely visual neurons. This calls for recasting of a hypothesized SC pathway for saccadic suppression, relying on a relay to superficial SC layers from deeper centers of the saccade motor command.

## Discussion

We found spatial-frequency selective saccadic suppression in SC visual-motor neurons, and the neural dynamics of visual-motor neuron suppression were well correlated with behavior. Our results are in line with interpretations of saccadic suppression as a reduction in response gain (Chen et al. 2015; Guez et al. 2013; Hafed and Krauzlis 2010). Consistent with this, we have recently found that neural contrast thresholds in the SC are altered around the time of microsaccades (Chen et al. 2015). We have also found that for SC neurons possessing some baseline activity in the absence of a visual stimulus, there was very modest peri-microsaccadic modulation of activity (see Fig. S2 of Chen et al. 2015) when compared to the modulations in stimulus-evoked visual bursts that we have observed here and earlier (Chen et al. 2015; Hafed and Krauzlis 2010). We believe that observations like these place constraints on the potential sources and mechanisms of extra-retinal modulation that is often invoked in theories of saccadic suppression.

There have been few successful demonstrations of perceptually-relevant patterns of saccadic suppression in neural activity. In the earliest visual areas, selective magno-cellular pathway suppression is not clear (Hass and Horwitz 2011; Kleiser et al. 2004; Ramcharan et al. 2001; Reppas et al. 2002; Royal et al. 2006), even though behavioral effects strongly predicted them (Burr et al. 1982; Burr et al. 1994; Hass and Horwitz 2011; Volkmann et al. 1978). Rather, there is mild suppression throughout these early areas, regardless of magno- or parvo-cellular pathway. Higher areas, primarily in the dorsal stream, do show saccadic suppression dynamics (Zanos et al. 2016; Bremmer et al. 2009; Han et al. 2009; Ibbotson et al. 2008; Ibbotson et al. 2007; Thiele et al. 2002), but the origins of such suppression remain elusive. In fact, it has been suggested that suppression in motion-related areas MT and MST (Bremmer et al. 2009; Ibbotson et al. 2008; Ibbotson et al. 2007; Thiele et al. 2002) may be inherited from earlier visual areas (Ibbotson et al. 2008; Ibbotson et al. 2007), which themselves have weak and unselective suppression. Thus, there is a pressing need for better understanding of saccadic suppression mechanisms.

The fact that primarily motion areas have been shown to exhibit the most convincing suppression additionally does not help account for the fact that saccadic suppression may be useful for perception even if the “motion problem” (Wurtz 2008) caused by saccades is solved. For example, suppression could help regularize processing of stimuli after saccades, regardless of the image shift itself. Consistent with this, we saw SC suppression for microsaccades, even though both the retinal-image motion and displacement caused by these eye movements are quite mild. Moreover, we saw suppression even with purely stationary gratings.

Related to the above, the fact that we saw any effects with microsaccades at all is interesting in its own regard, but the real advantage from studying microsaccades was that they allowed better experimental control. Microsaccades are mechanistically similar to larger saccades (Hafed 2011; Hafed et al. 2015; Hafed et al. 2009; Zuber et al. 1965), making them an extremely viable tool to understanding saccadic suppression. However, these movements simplify several challenges associated with large saccades. For example, studies with large saccades have to contend with large image shifts caused by eye movements. As a result, full field stimuli become necessary (Ibbotson et al. 2008; Ibbotson et al. 2007). However, in our case, we could use stimuli identical to how normal experiments might stimulate RF’s.

Another experimental advantage here was the fact that SC shows suppression *after* saccades in our type of paradigm (Chen et al. 2015; Hafed and Krauzlis 2010). This allowed us to avoid probing neurons during the eye movements themselves; we always presented stimuli *after* microsaccades, such that no-microsaccade and microsaccade trials both had the exact same stimulus, location, and eye-state. Of course, saccadic suppression in the SC would be even stronger *during* the microsaccades themselves, as we have recently shown (Chen et al. 2015; Hafed and Krauzlis 2010), which is further evidence of a consistency between our visual-motor neural modulations and classic perceptual effects of saccadic suppression in humans. That is, our choice to focus in this paper on post-movement modulations was one of exploiting the experimental advantages of doing so as opposed to one of a conceptual difference between our visual-motor neural modulations and the perceptual phenomenon itself.

Concerning superficial visual neurons, one question arises on the sources of mild and unselective suppression that we saw. This could reflect retinal effects. For example, retinal outputs show transient perturbations in response to saccade-like image displacements (Roska and Werblin 2003). Additionally, the effect could be inherited from V1, which also does not show selectivity (Hass and Horwitz 2011). Regardless of the source, what is clear is that suppression in visual neurons is not selective as in perception. However, it could still be functional. For example, a collicular-cortical pathway from superficial SC may selectively target motion-related areas (Berman and Wurtz 2008; 2010; 2011; Wurtz et al. 2011). As a result, superficial SC may still contribute to saccadic suppression of motion (Bridgeman et al. 1975; Burr et al. 1982); in this case, selectively suppressing motion by superficial SC neurons would arise not necessarily because the neurons themselves are selective in their suppression profiles, but instead because of selectivity in their connections to cortical targets.

Our observation of a lack of suppression selectivity in purely visual neurons also helps address an important question regarding the nature of our selective visual-motor neuron modulations. Specifically, it may be argued that (peripheral) SC neurons may preferentially over-sample low spatial frequencies in their tuning curves (Hafed and Chen 2016), meaning that they exhibit higher baseline sensitivity for low spatial frequencies even without microsaccades. This, in turn, could mean that we only saw stronger suppression at low spatial frequencies (in the visual-motor neurons) simply because the baseline visual responses were stronger; suppression could in reality be constant across spatial frequencies, but its effect on absolute firing rate would scale with visual response sensitivity. However, in our mappings of SC tuning curves, we found that purely visual neurons, like visual-motor neurons, also tend to be more sensitive to low spatial frequencies. If our effects are explained by the dependence of suppression on baseline visual sensitivity in the absence of microsaccades, then our purely visual neurons should have shown the same patterns of selective suppression as the visual-motor neurons. They did not (Figs. 3, 4). Second, we specifically examined suppression within each spatial frequency relative to the no-microsaccade baseline of the same frequency, in order to isolate the suppression effect independent of baseline response strength. This avoided questions of absolute firing sensitivity across spatial frequencies. Third, in Fig. 7, we explicitly examined suppression as a function of preferred spatial frequency and still found diminishing returns in suppression strength with increasing spatial frequency even when each spatial frequency bin only included the neurons preferring that frequency.

Finally, because the visual system is inherently generally low pass anyway (especially in the periphery), then even a mechanism in which suppression simply scales with visual sensitivity of a given spatial frequency would still explain the well known perceptual phenomenon of selective suppression of low spatial frequencies in humans.

There may also be an additional potential counter-interpretation of our results. Specifically, it may be argued that we uncovered a highly specific effect only modulating saccadic RT’s, and that visual-motor neuron modulations are irrelevant for other forms of behavior (e.g. not requiring saccadic responses). However, this is highly unlikely. First, the SC contributes to behavior even with non-saccadic outputs. For example, during attentional tasks with button presses, SC lesions impair performance (Sapir et al. 1999), suggesting that it is sensory and/or cognitive modulations that are relevant. Consistent with this, the SC contributes to attentional paradigms with a variety of response modalities (Lovejoy and Krauzlis 2010; Zenon and Krauzlis 2012). Second, we only looked at the earliest visual responses and uncovered strong correlations to behavior observed in separate experiments. This indicates that it was the *sensory response* that mattered. Consistent with this, we have recently found that microsaccades have a virtually identical impact on either saccadic or manual responses (Tian et al. 2016). Third, our behavioral effects on RT are themselves remarkably similar to perceptual effects of saccadic suppression in humans, but using different perceptual measures and response modalities (Burr et al. 1982; Burr et al. 1994; Volkmann et al. 1978). Fourth, we found that monkey P had a stronger suppression effect in behavior than monkey N at the low spatial frequencies (compare the cyan curves in Fig. 2C and Fig. 2F) even though monkey P had significantly longer saccadic RT’s to begin with (compare the black no-microsaccade curves in Fig. 2A and Fig. 2D). If our behavioral and neural effects were restricted to limits on saccadic RT, perhaps due to potential saccadic refractory periods between successive saccades and microsaccades, then monkey P should have shown weaker behavioral suppression than monkey N since this monkey’s saccadic system had plenty of time to recover from the previous generation of a microsaccade before having to generate the next saccadic RT. Given all of the above, we find it highly unlikely that our modulations are only specific to modulating saccadic RT’s.

Given the above, it might be additionally asked why the SC should be among the neural substrates for perceptually-relevant saccadic suppression? We think that the SC has several appealing features to place it well within a hypothetical saccadic suppression system. For example, the SC contributes to triggering the saccade command. Thus, a source of corollary discharge is already present in the visual-motor layers, as demonstrated by our differential firing rate (Figs. 3-7) and LFP (Figs. 8-10) effects.

Second, proximity of the SC to motor outputs confers an additional advantage: SC suppression, besides having perceptual effects, could help to regularize how often subsequent saccades are made to sample the visual world. That is, in reality, suppression could serve to control the temporal structure of saccades, which can be very important both behaviorally (Tian et al. 2016) and cortically (Lowet et al. 2016). Finally, our results are in line with an early hypothesis that the primary output of SC may be the visual-motor layers (Mohler and Wurtz 1976).

In all, our results will motivate further investigation of a classic, yet highly controversial topic in systems neuroscience.

## Acknowledgements

We were funded by the Werner Reichardt Centre for Integrative Neuroscience (CIN) at the Eberhard Karls University of Tübingen. The CIN is an Excellence Cluster funded by the Deutsche Forschungsgemeinschaft (DFG) within the framework of the Excellence Initiative (EXC 307).

